# DIMEdb: an integrated database and web service for metabolite identification in direct infusion mass spectrometery

**DOI:** 10.1101/291799

**Authors:** Keiron O’Shea, Divya Kattupalli, Luis AJ Mur, Nigel W Hardy, Biswapriya B Misra, Chuan Lu

## Abstract

**Motivation:** Metabolomics involves the characterisation, identification, and quantification of small molecules (metabolites) that act as the reaction intermediates of biological processes. Over the past few years, we have seen wide scale improvements in data processing, database, and statistical analysis tools. Direct infusion mass spectrometery (DIMS) is a widely used platform that is able to produce a global fingerprint of the metabolome, without the requirement of a prior chromatographic step - making it ideal for wide scale high-throughput metabolomics analysis. In spite of these developments, metabolite identification still remains a key bottleneck in untargeted mass spectrometry-based metabolomics studies. The first step of the metabolite identification task is to query masses against a metaboite database to get putative metabolite annotations. Each existing metabolite database differs in a number of aspects including coverage, format, and accessibility - often limiting the user to a rudimentary web interface. Manually combining multiple search results for a single experiment where there may be potentially hundreds of masses to investigate becomes an incredibly arduous task.

**Results:** To facilitate unified access to metabolite information we have created the Direct Infusion MEtabolite database (DIMEdb), a comprehensive web-based metabolite database that contains over 80,000 metabolites sourced from a number of renowned metabolite databases of which can be utilised in the analysis and annotation of DIMS data. To demostrate the efficacy of DIMEdb, a simple use case for metabolic identification is presented. DIMEdb aims to provide a single point of access to metabolite information, and hopefully facilitate the development of much needed bioinformatic tools.

**Availability:** DIMEdb is freely available at https://dimedb.ibers.aber.ac.uk.

**Supplementary information:** Supplementary data are available at *Bioinformatics* online.

## 1 Introduction

Metabolomics is the global study of naturally occuring low molecular weight organic molecules (metabolites) within cell, tissue, organ, or biofluid (German *et al.*, 2005) - providing a ‘snapshot’ of an organism’s metabolic state. The analysis of numerous metabolites is a growing area of analytical biochemistry. Metabolites act as the reaction intermediates of multiple biological processes, and are considered to be the key indicator of biochemical activities within an organism (Schmidt, 2004). Therefore metabolomics is widely used to a number of fields including plant science, toxicology, food science and biomedical research (O’Shea *et al.*, 2016; Stewart and Bolt, 2011; Mamas *et al.*, 2011).

Mass-spectrometry (MS)-based metabolomics studies are often separated into three main categories: targeted (Weljie *et al.*, 2006), semi-targeted (Burgess *et al.*, 2011), and untargeted analysis (De Vos *et al.*, 2007). Each strategy differs in multiple aspects including the quantification method (relative or absolute), sample preparation protocols, spectral accuracy, spectral resolution, the total number of metabolites detected, and even the overall study objective. In targeted and semi-targeted analysis the metabolite composition of the studied material is known prior to any experimentation. Whilst this approach greatly simplifies the metabolite identification process, there is a limit to the number of commercially available standards, greatly inhibiting the potentiality of scientifically interesting novel discoveries. Untargeted analysis, commonly referred to as metabolite ‘fingerprinting’, describes the study of the global metabolome of the studied species. Whilst this approach often results in the most diverse and comprehensive snapshot of the studied metabolome, it greatly inhibits accurate metabolite identification.

There exists an array of analytical platforms, suited for metabolomic analysis, however the two technologies synonymous with the field are mass spectroscopy (MS) (Dettmer *et al.*, 2007) and nuclear magnetic resonance (NMR) (Lin *et al.*, 2007) - both of which are capable of producing high dimensional high resolution quantitative metabolite data. MS utilises the formation of positive and/or negatively charged ions from a given metabolite in the sample before ion separation in accordance to mass-to-charge ratios (m/z). MS platforms including gas-chromatography mass spectroscopy (GC-MS) (Kanani *et al.*, 2008)), liquid chromatography (LC-MS) (De Vos *et al.*, 2007) and their ultra-high performance (UHP) successors (Roux *et al.*, 2012) are the most prevalent platforms for the global analysis of the metabolome. Despite their popularity, both platforms suffer from their own restrictions. For example, in GC-MS only volatile metabolites are quantified. LC-MS is known to show inconsistencies in peak quality and retention time - resulting in poor peak alignment and misalignment and leading to a potentially misreported analysis. Moreover, the requirement of prior chromatographic separation before analysis makes performing large-scale high-throughput MS assays using these technologies a time-consuming and physically exhaustive process. Recently the use of direct infusion mass spectroscopy (DIMS) has become a popular platform for high-throughput metabolomic analyses (Koulman *et al.*, 2007). DIMS skips the chromatographic step in its entirety, instead allow for samples to be directly introduced to the mass spectrometer - a task which can be automated through the use of flow injection technologies.

The identification of even a few metabolites within spectrum produced from untargeted mass spectrometry analysis is both incredibly challenging and labour intensive (Moco *et al.*, 2007), and despite vast advancement in analysis technology and protocols still remains a major bottleneck to users (Dunn *et al.*, 2013). When attempting to identify metabolites found within metabolomic samples, consideration needs to be made for only the differences between metabolites of different nominal mass - but more often than not between metabolites with the same nominal mass yet dissimilar molecular composition. Moreover, MS-based metabolomics detects metabolites as a collection of derived species, and thus correctly identifying the peak (or parent) metabolite is essential for accurate identification. Another major issue facing the identification task is that the complete composition of the metabolome remains unknown (Nakabayashi and Saito, 2013) - making the unambiguous identification of metaboloties within untargeted mass spectrum without the use of further chemical analysis unachievable in many cases. In accordance to guidelines set out by the Metabolomics Standards Initiative (MSI), the identification of a metabolite at the highest confidence level (1) requires comparisons of a species orthogonal properties against an authentic standard measured with the exact same experimental conditions. Unfortunately due to the lack of available authentic compounds, this form of metabolite identification may be impossible to achieve in many cases (Sumner *et al.*, 2007).

There exists a number of online metabolomics databases that can be utilised for the identification of metabolites from a given mass spectra. These databases often differ in their coverage, with many being species or organism-specific (e.g. Human Metabolite Database (HMDB) (Wishart *et al.*, 2007), Seaweed Metabolite Database (SWMD) (Davis and Vasanthi, 2011), Metabolome Tomato Database (MoToDB) (Moco *et al.*, 2006)), or only suited for a specific analytic tool (i.e, for GC-MS; GOLM Hummel *et al.* (2007)). Moreover, existing databases that have attempted to collate an extensive integrative DIMS-compliant platform are no longer regulary maintained or updated (e.g. MZedDB Draper *et al.* (2009)). As it is desirable to perform mass search over multiple datasets (as to both confirm results), researchers are having to manually perform this task ask each metabolite spectral resource limits its data through only a web application. Hence, there is a need for both user-friendly and programmatic access to a well curated up-to-date metabolite database.

To this end, we have developed the Direct Infusion MEtabolomics database (DIMEdb), which provides both programmatic and user friendly interfaces to an integrated metabolite database - sourced from multiple popular metabolomic and chemical databases. In addition to providing a web-based front end to metabolite data, we hope to enable bioinformaticians to build and extend much needed bioinformatic tools for the analysis and integration of metabolite-related data.

## 2 Methods

### 2.1 Database construction

We chose a number of highly populated metabolite databases including HMDB, PubChem, CHEBI, BioCyc, and KEGG for integration. As each of these databases came in unique formats they were combined into a standard form containing key attributes including naming conventions, unique identifiers (namely Simplified Molecular Input Line Entry System representation (SMILES) and/or IUPAC International Chemical Identifier (InChI) where available), and physciochemical information. Upon inspection, we found that physciochemical information found throughout these databases was prone to a high level of variability. Moreover, chemical databases (such as PubChem and BioCyc) presented information as a charged molecule. As a result of this, an in-silico approach (using the RDKit (Landrum, 2006) cheminformatics toolkit) was used to calculate the physicochemical for each given metabolite. Once physicochemical properties were calculated, molecule entries were combined through the use of a generated InChIKey - a 25 character hashed representation of the InChI identifier. In instances where unexpected collisions were present (e.g. the presence of corrupted hashed keys), the BridgeDb identification mapping service (van Iersel *et al.*, 2010) was used to match entries in accordance to external resources (including KEGG, HMDB, BioCyc, MetaCyc). As to filter out redundant compounds, entries with less than 6 elements were removed.

As metabolites exist within a biological context, integration of metabolite networks and biological pathway information was vital in enhancing the metabolite identification process. Whilst some collated databases contained pathway/network information, most databases did not. Using mapped metabolites and their given accessions, we were able to make use of web services provided by BioCyc (Caspi *et al.*, 2007), KEGG (Kanehisa and Goto, 2000), and SMPDB (Frolkis *et al.*, 2009) to retrieve pathway information and integreate contextual information into our database.

As noted earlier, in-source fragmentation of molecules are able to provide key structural information that can be used for metabolite identification. Using adjusted molecular formulae, it is possible to calculate accurate masses given likely ionisation rules (Keller *et al.*, 2008). Unfortunately existing *in-silico* molecular weight calculators are closed off for programmatic use, making them impractical for database composition. The Python Isotopic Distribution Calculator (O’Shea, 2017) was written and released to calculate mass spectrometry adducts.

### 2.2 Interface

To facilitate greater extensibility, every possible effort has been made to ensure that DIMEdb’s underlying architecture is as modular as possible. DIMEdb can be broken down into two main facets; a Representational State Transfer Application Programming Interface (RESTful API), and a web application that makes use of the API to query and interface the generated metabolites a user-friendly web interface (Figure 1).

**Fig. 1.**
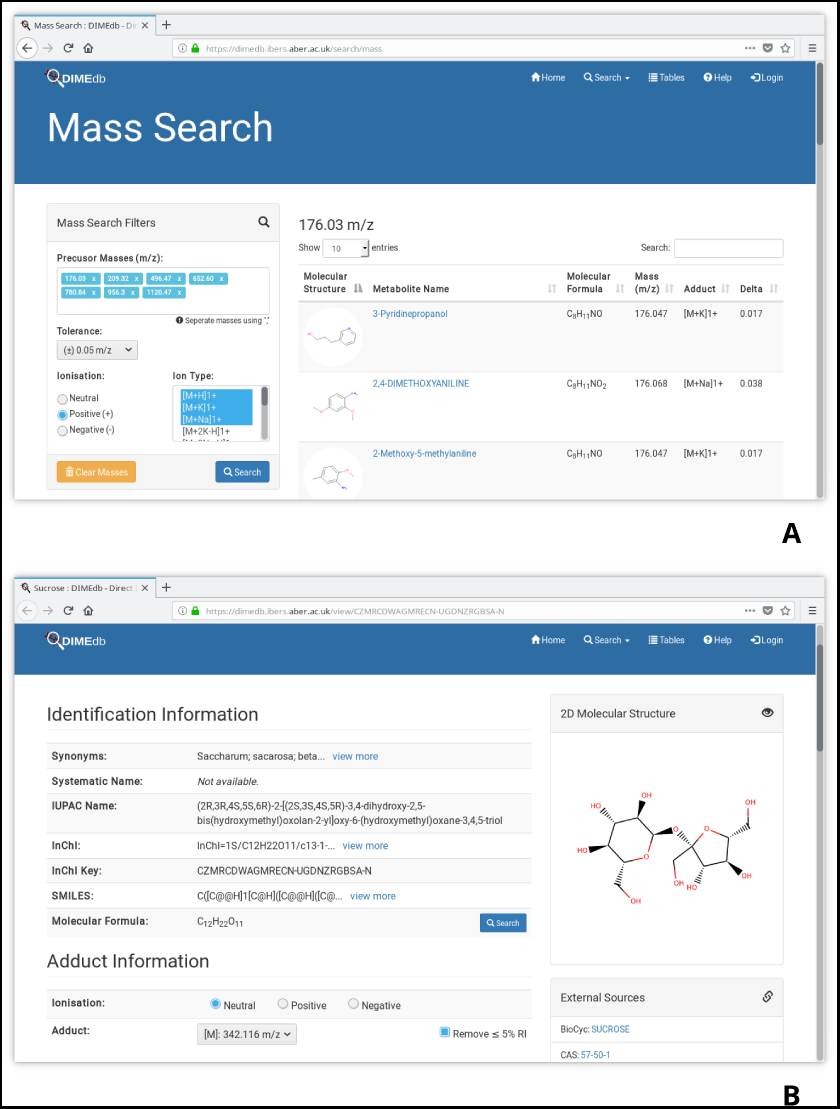
Screenshots of the DIMEdb web interface. Panel A shows results for a list of mass queries. Panel B provides detailed metabolite information, including identification information, adduct information, a 2D visual representation, external resources, and contextual information for sucrose.

**Fig. 2.**
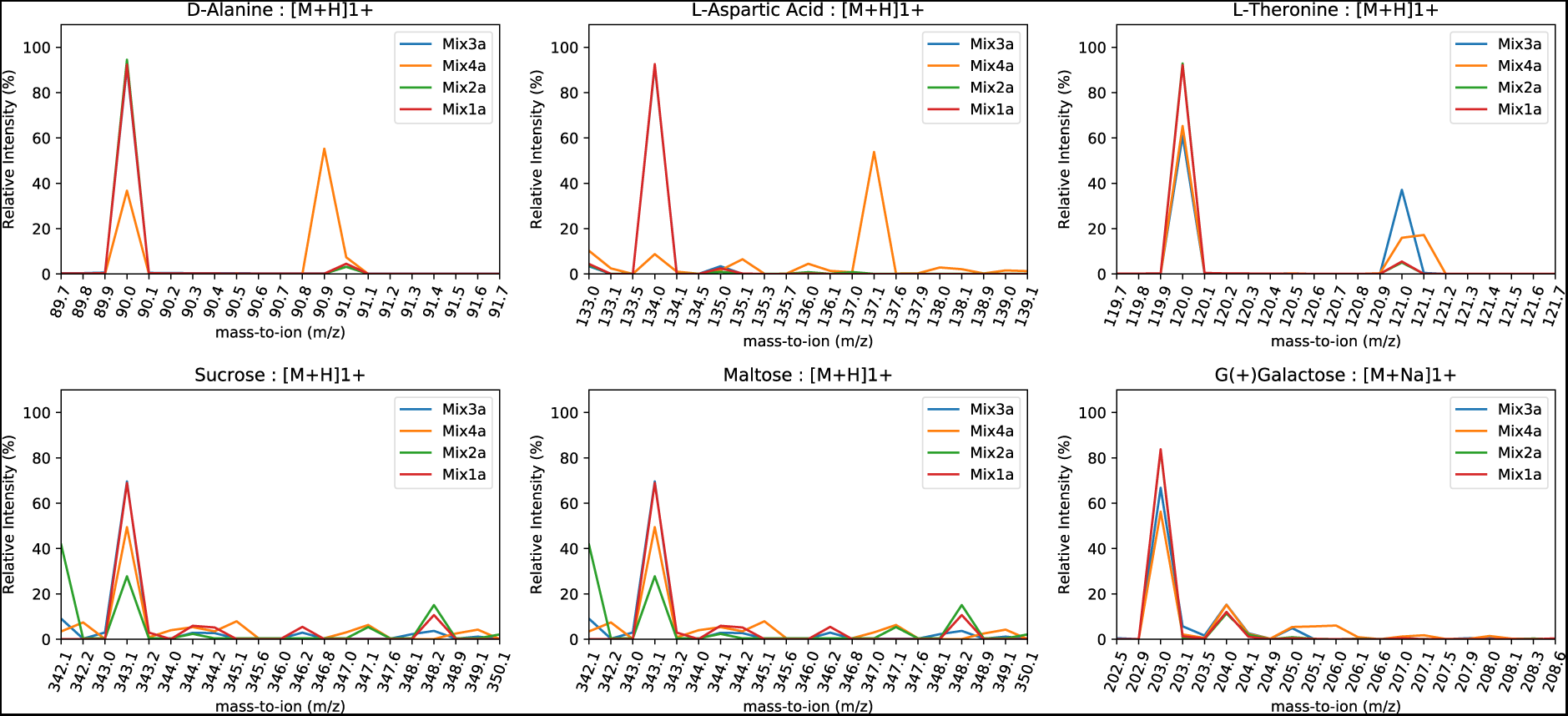
Mix-averaged Flow-Infusion Electrospray Mass Spectrometry (FIE-MS) plots of identified metabolites in positive ion mode.

The web interface provides two different methods of searching the metabolite database; a text-based search and a precursor mass search. The text-base search page facilitates users to search the metabolite database by searching by a number of parameters-including the systematic chemical name, chemical nomenclature provided through literature and existing databases, database accessions (including HMDB, ChEBI, PubChem, KEGG, CAS, and Wikidata), InChI, InChI Key, SMILES and molecular formula. The results pane displays a table containing all metabolites matching the user specified search criteria. The results table contains a 2D graphical representation of the metabolite, metabolite name, molecular formula, and monoisotopic weight.

In addition to text search, mass-based search facilities are provided which allows for metabolite identification from a batch of precursor mass-to-ions. The user may also choose to set parameters such as adducts and a mass tolerance, as to better limit the search space. Using this tool, users are able to determine the molecular formula and potential chemical annotations for a given batch of mass values.

RESTful APIs provide an interface via which programs and web services can query and retrieve information that corresponds to the user’s request. Queries can be sent through an HTTP POST request to the URL that specifies a given resource, which will instruct the server to produce information of interest. DIMEdb’s web service is detailed on the online documentation, empowering users to create web service clients that can make use of our comprehensive metabolite database for their own applications. To demonstrate this functionality, an example Python client has been written and can be found in the aforementioned DIMEdb documentation.

### 2.3 Implementation

DIMEdb is built on the Python Flask (http://flask.pocoo.org/) web application microframework. Metabolite information is stored in the document-orientated MongoDB database. User account information is stored in a relational Postgresql database, and encrypted using the bcrypt Python package. Using a high performance document-orientated database enables us to provide a substantial database on a low-cost virtual machine without taking a substantial performance decrease when indexing the database. The DIMEdb RESTful API was implemented using the Python Eve (http://python-eve.org/) framework, with custom interfaces written using PyMongo (https://api.mongodb.com/python/current/). The web interface was written in the Jinja templating engine, using Bootstrap and numerous JavaScript libraries to enable a rich user interface.

## 3 Results and Discussion

To demonstrate the effectiveness of the DIMEdb platform, solutions of sugars (sucrose, maltose, galactose), amino acids (alanine, threonine, aspartic acid) and fatty acids (palmitic acid, stearic acid, pentadecanoic acid) were prepared in 75% methanol/water (v/v) for identification purposes. Cocktails of four mixes were made with all the compounds but at different concentrations in each mix. In the sample mix A, sucrose (10mM) is the highest whereas pentadecanoic acid (0.08mM) is the lowest concentrated compounds. Mix B contains more concentration of threonine (10mM) and less of palmitic acid (0.08mM). The Mix C is made up of increased concentration of sucrose (5mM) and decreased stearic acid (0.03mM). In mix D, threonine (0.5mM) is in excess and pentadecanoic acid (0.003mM) is low compared to all other compounds. A master mix is made by blending all the 4 sample mixes, and 75% methanol is used as a blank. The randomised sample injections were preceded for metabolite profiling by flow infusion into a Thermo Scientific^TM^ Exactive^TM^ Plus Orbitrap Mass Spectrometer. The Theano RAW data files were converted into the open standard mzML file format (Martens *et al.*, 2011), before being processed.

### 3.1 Metabolomics data processing

Metabolomic processing steps detailed have been reported as suggested by the Metabolomics Standards Initiative (**?**) using the dimspy package (O’Shea, 2018). Whereas other forms of MS produces multiple ‘scans’ to be aligned into a single spectrum (feature matrix) through the use of spectral alignment tools, DIMS produces a single summed (or averaged) matrix consisting only of masses and intensities -disposing of the retention time property altogether (Castrillo *et al.*, 2003). Generated mzML files were analysed by their total ion count (TIC) to check for any unusual samples (where ionisation had failed, or an instrumental error had occurred) - from which no samples were found to be erroneous. As to improve the signal quality and reduce artifacts, several pre-processing steps were applied. Adaptive iteratively reweighted Penalized Least Squares (airPLS) baseline correction (Zhang and Liang, 2012) was employed to remove low frequency ‘noisy’ artifacts that were produced by experimental and instrumental conditions.

Baseline corrected data was then subjected to peak detection and alignment, in which peaks were identified using a Gaussian wave filter, where a Gaussian filter is applied to the spectrum before being centroided to locate peaks within the spectrum (Smith *et al.*, 2006). Standard metabolomic-analysis protocols make use of statistical methods to determine masses of interest. As we had no phenotype criteria of which to highlight affecting metabolites, we resorted to simply comparing the mix spectra to clean (empty) runs to locate the presence of any substantial peak differences. The peaks with the largest log-fold change between the samples were taken forward for metabolite identification.

### 3.2 Metabolite identification with isotopic distribution matching

Even with advent of ultra-high resolution mass spectrometers, it is not always possible to accurately identify metabolites using just the mass value. As to to reduce the enormity of the mass-search space, it is important to consider a number of molecular properties. Data produced from DIMS produces thousands of metabolite features, where a single metabolite may be detected as multiple separate features (including protonated and deprotonated ions, adducts, and isotopomers). Isotope identification allows us to significantly scale down the search-space through limiting the potential adduct relationship found for a given mass.

There exists a number of computational methodologies for the identification of metabolites using adduct and isotopic pattern information (Uppal *et al.*, 2017; Brown *et al.*, 2009; Silva *et al.*, 2014). However, we have opted to use a commonly used pairwise correlation analysis-based approach for ion and isotope identification. Pearson’s pairwise correlation analysis of all the masses in the processed feature matrix was calculated using the Pandas data analysis package (McKinney, 2011). This produced a matrix of correlation coefficients for pairs of masses, which indicates the likelihood of mass pairs coming from the products of the same metablilte. Only feature pairs of which showed a coefficient score of *≥* 0.9 correlation values were kept for further analysis.

Agglomorative clustering were performed using the complete linkage algorithm to locate clusters of correlating masses, from which a dendrogram was produced to help visualise the identified relationship clusters-highlight potential presence of an isotope pattern. Using one of the key mass signals that was shown to be a clear differentiator between clean and mix sample (203 *m/z*), two additional signals were found to have been highly correlated (204 and 205 *m/z*). On further manual inspection, the isotopic pattern was shown to match that of a Na-adduct-with 203 *m/z* being the peak abundance.

Using this information a simple paramaterised mass search was conducted using the DIMEdb web interface, limiting the search to +/-0.25 *m/z* of 203.053 *m/z* (the averaged accurate mass found within the peak bin) of all positivie charged metabolite entries featuring a [M+Na]1+ ion type. This returned a total of 4 metabolite entries, of which all had the same nominal chemical formula (C_6_H_12_O_6_)-corresponding to a monosaccharide galactose. This process was repeated in both positive and negative mode ionisation mode spectrum (results can be found in Table 1).

**Table 1.**
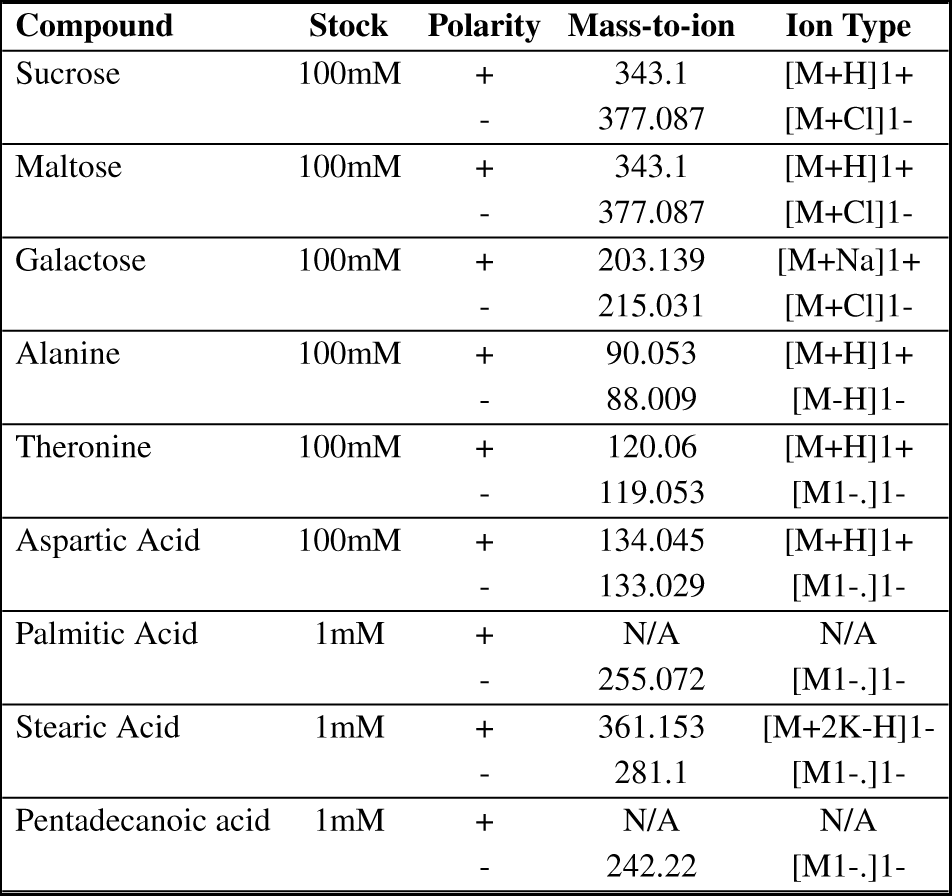
Metabolite annotation table of mass values identified from both positive and negatively charged FIE-MS. As expected smaller quantities of fatty acids did not show up in positive polarity mass spectrum.

## 4 Conclusions and Future Work

There is an abundance of chemical and metabolite information readily available in the public domain. Unfortunately, making use of this vast array of information is difficult due to a lack of standard formatting and data interrogation techniques. We have developed DIMEdb with the goal of centeralising multiple renowned metabolite databases into an easily accessible and highly functional web service. We have curated thousands of metabolites, and through our powerful data generator have been able to assure a high level of data quality. We are continuously releasing revised versions of the database with both new and curated data, and hope to facilitate user driven revisions in the near future. Moreover, it is our goal to further enhance contextual analysis of metabolites through the incorporation of pathway and metabolite network analysis tools.

DIMEdb is freely available to download and use under the open source MIT license, and can be freely accessed at https://dimedb.ibers.aber.ac.uk. Full documentation and additional software packages can be located at http://dimedb.ibers.aber.ac.uk/help.

